# OXSR1 inhibits inflammasome activation by limiting potassium efflux during mycobacterial infection

**DOI:** 10.1101/2021.04.21.440692

**Authors:** Elinor Hortle, Lam Vi Tran, Angela RM Fontaine, Natalia Pinello, Justin J-L Wong, Warwick J Britton, Stefan H Oehlers

## Abstract

Pathogenic mycobacteria inhibit inflammasome activation as part of their pathogenesis. While it is known that potassium efflux is a trigger for inflammasome activation, the interaction between mycobacterial infection, potassium efflux and inflammasome activation has not been investigated. Here we use *Mycobacterium marinum* infection of zebrafish embryos and *Mycobacterium tuberculosis* of human THP-1 cells to demonstrate that pathogenic mycobacteria upregulate the host WNK signalling pathway kinases SPAK and OXSR1 which control intracellular potassium balance. We show that genetic depletion or inhibition of OXSR1 decreases bacterial burden and intracellular potassium levels. The protective effects of OXSR1 depletion are mediated by NLRP3 inflammasome activation and are dependent on caspase-mediated release of IL-1β and the downstream activation of protective TNF-α. The elucidation of this druggable pathway to potentiate inflammasome activation provides a new avenue for the development of host-directed therapies against intracellular infections.

## Introduction

Inflammasomes are large cytosolic multi-protein complexes that are critical for the immune response to infection. They facilitate the production of bioactive IL-1β^1^, which directs pathogen killing through upregulation of TNF-α^2^ and orchestrates systemic immune control through paracrine signalling. Inflammasomes typically consist of a sensor protein, which detects specific stimuli within the cytosol, and an adaptor protein which facilitates the oligomerization of the sensor with pro-caspase-1 ^3,4^. Inflammasome assembly triggers activation of caspase-1, which then cleaves pro-IL-1β and pro-IL-18 into their active forms. Caspase-1 also cleaves Gasdermin D, the N-terminal fragment of which forms pores in the cell membrane, allowing secretion of active IL-1β and IL-18, and triggering cell death via pyroptosis ^4^. These events contribute to host defence by both rapidly inducing the inflammatory response and limiting replication of intracellular pathogens.

To escape this immune control, many successful intracellular pathogens have evolved methods to limit inflammasome activation ^3^. Influenza A, *Pseudomonas aeruginosa*, Baculovirus, Vaccinia virus, *Streptococcus pneumoniae*, Myxoma virus and *Yersinia pseudotuberculosis* have all been shown to limit IL-1β production by inhibiting caspase-1 activation^3^. In the case of pathogenic mycobacteria, their interactions with inflammasome activation are more complex. One study has shown that *Mycobacterium tuberculosis* actively inhibits inflammasome activation via a zinc metalloprotease^5^, and that clinical isolates associated with severe disease evade NLRP3 activation ^6^. Others have shown that *M. tuberculosis* induces both NLRP3 inflammasome and caspase-1 activation ^7-9^. Further studies suggest that *M. tuberculosis* actively inhibits activation of the AIM2 inflammasome and dampens activation of the NLRP3 inflammasome by upregulation of NOS and IFN-β^1,9-12^.

The NLRP3 inflammasome, one of the best studied inflammasomes, can be triggered by numerous stimuli, including ATP, heme, pathogen-associated RNA, and a variety of bacterial components^13-17^. Because these triggers are so diverse, it has been suspected that these activation stimuli are not detected by NLRP3 directly, but rather NLRP3 activation is the result of converging cellular signals. There is evidence that mitochondrial dysfunction, reactive oxygen species, and lysosomal damage contribute to NLRP3 activation (as reviewed elsewhere ^18,19^). A common event that occurs downstream of almost every NLRP3 stimulus is potassium (K^+^) efflux. Studies have shown that K^+^ ionophores stimulate NLRP3^20^, that high extracellular K^+^ can inhibit NLRP3 activation^21,22^, and that K^+^ efflux alone is sufficient to activate the NLRP3 inflammasome^23^. Although the mechanism linking K^+^ efflux to activation of NLRP3 is not well defined, evidence suggests that K^+^ efflux occurs upstream of NLRP3 activation and may induce a conformational change in NLRP3 that favours oligemerization^23,24^. This raises the possibility that K^+^ efflux pathways could be targeted to therapeutically activate, or potentiate the activation of, NLRP3 to control intracellular pathogens.

One of the master regulators of cellular K^+^ flux is the With-No-Lysine (WNK) kinase signalling pathway. In response to cellular stress or osmotic changes, WNK kinase activates the SPAK and OXSR1 kinases. SPAK and OXSR1 inhibit the KCC channels, which pump K^+^ out of the cell, and activate the NKCC channels, which pump K^+^ into the cell^25^. It has previously been shown that blocking the interaction of SPAK/OXSR1 and KCC1 leads to net K^+^ efflux from the cell^26^. We have further shown that constitutively active KCC1 alters the inflammatory response to malaria infection in mice, and that this effect is associated with dramatically increased survival^27^. Here we sought to investigate whether the SPAK/OXSR1 pathway is involved in the host response to mycobacterial infection, and if this pathway could be manipulated as a host-directed therapy against infection.

## Results

### Infection-induced activation of *oxsr1a* aids the growth of pathogenic mycobacteria

To determine if SPAK and OXSR1 are involved in immunity, we infected zebrafish embryos with *Mycobacterium marinum* and analysed gene expression at 3 days post infection (dpi). Both *stk39* and *oxsr1a* (the zebrafish orthologs of *SPAK* and *OXSR1* respectively) were significantly upregulated at 3 dpi compared to uninfected embryos (Figure 1A). This result is consistent with previous data showing *oxsr1a* is upregulated in *M. marinum*-infected macrophages^28^.

**Figure 1:**
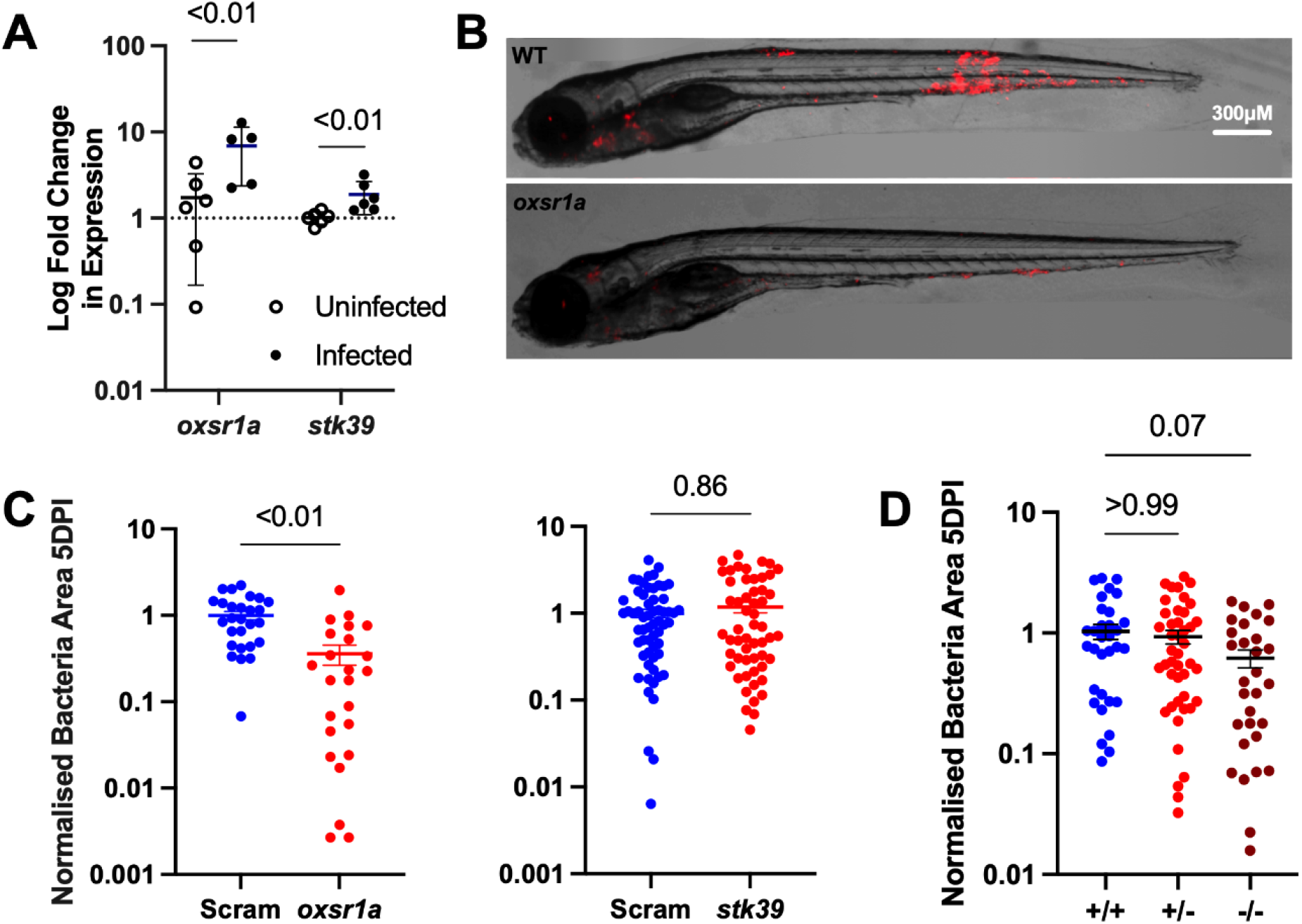
Infection-induced OXSR1 aids the growth of *M. marinum* in zebrafish. A) Relative expression of *oxsr1a* and *stk39* in 3 dpi *M. marinum*-infected zebrafish embryos, compared to matched uninfected controls. Biological replicates (n=6) each represent pooled RNA from 7-10 embryos. B) Representative images of *M*. marinum-tdTomato (red) bacterial burden in scramble control and mosaic F0 *oxsr1a* crispant embryos. C) Quantification of *M. marinum* bacterial burden in scramble control, mosaic F0 *oxsr1a* and *stk39* crispant embryos. Each graph shows combined results of two independent experiments. D) Quantification of *M. marinum* bacterial burden in WT, heterozygous and homozygous *oxsr1a* knockout embryos. Graph shows combined results of two independent experiments.

To determine if the upregulation of *stk39* and *oxsr1a* results in increased bacterial growth, we depleted each kinase individually by CRISPR-Cas9 knockdown and infected the embryos with fluorescent *M. marinum* (Extended Data 1). *M. marinum* burden was significantly reduced in *oxsr1a*, but not *stk39*, knockdown embryos (Figures 1B and 1C). To confirm these results, we created a stable *oxsr1a* knockout allele *oxsr1a*^*syd5*^(Extended Data 2). Homozygous, but not heterozygous, *oxsr1a*^*syd5*^ embryos showed reduced bacterial burden (Figure 1D).

### The immunomodulatory role of OXSR1 is conserved in human cells

To determine whether the immunomodulatory role of OXSR1 is conserved across species, we first differentiated human THP-1 cells with PMA and infected with *M. tuberculosis*. At 3 dpi OXSR1 protein expression was significantly upregulated in infected cells compared to uninfected cells (Figure 2A and Extended Data 3), mirroring the increased *oxsr1a* expression observed in infected zebrafish embryos. To determine whether this upregulation would affect bacterial burden, we generated an *OXSR1* knockdown human THP-1 cell line (Figures 2B and Extended Data 3).

**Figure 2:**
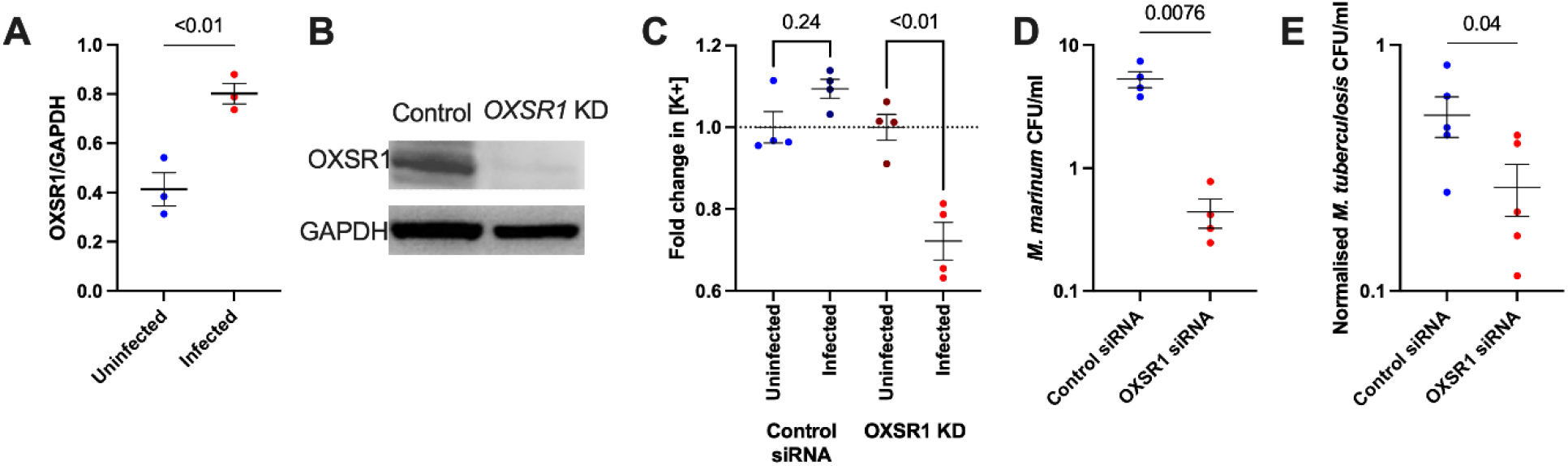
OXSR1 aids the growth of *M. tuberculosis* in human THP-1 cells. A) Quantification of OXSR1 protein in *M. tuberculosis*-infected PMA-differentiated THP-1 cells. Blots are included in Extended data 3. B) Western blot of *OXSR1* knockdown and vector control THP-1 cell lines, showing loss of OXSR1. Full un-edited blots are included in Extended data 3. C) Fold change of ION K+ green mean fluorescence intensity (measured by flow cytometry) of *M. marinum* infected control and *OXSR1* knockdown THP-1 cells. Fold change was calculated by dividing the MFI of cells positive for *M. marinum*-Katushka by the MFI of cells from the uninfected group. D) Quantification of intracellular *M. marinum* burden in 1 dpi *OXSR1* knockdown PMA-differentiated THP-1 cells. E) Quantification of intracellular *M. tuberculosis* burden in 3 dpi *OXSR1* knockdown PMA-differentiated THP-1 cells.

We first differentiated these THP-1 knockdown cells with PMA and infected with *M. marinum* to determine if mycobacterial infection would induce changes in intracellular K^+^ concentration. While we saw a small, statistically insignificant, increase in intracellular K^+^ concentration in *M. marinum*-infected control siRNA-expressing cells, *M. marinum*-infected *OXSR1* knockdown cells had significantly reduced K^+^ concentration compared to uninfected *OXSR1* knockdown cells (Figure 2C).

We next infected our *OXSR1* knockdown THP-1 cells with *M. marinum* and *M. tuberculosis* H37Rv and quantified bacterial growth by CFU recovery. At 1 dpi, knockdown THP-1 cells had reduced intracellular *M. marinum* load compared to control THP-1 cells (Figure 2D). At 3 dpi, knockdown THP-1 cells had reduced intracellular *M. tuberculosis* load compared to WT THP-1 cells (Figure 2E).

Together these results indicate that OXSR1 controls cellular potassium flux during mycobacterial infection and that the immunomodulatory role of OXSR1 is conserved between zebrafish and humans.

### Small molecule inhibition of SPAK/OXSR1 is host protective

Because SPAK/OXSR1-modulated K^+^ channels also shuttle Cl^-^, Na^2+^ and Ca^2+^, this pathway has been studied for its role in hypertension. The small molecule, Compound B (CB), reduces hypertension in animal models by inhibiting WNK phosphorylation of SPAK/OXSR1^29^, preventing SPAK/OXSR1 activation^30^. We first determined that 1.8 μM was the maximum dose of CB that could be tolerated by zebrafish larvae for 5 days for infection (Table 4). This concentration of CB did not affect the growth of *M. marinum* in axenic culture (Figure 3A).

**Table 1.**
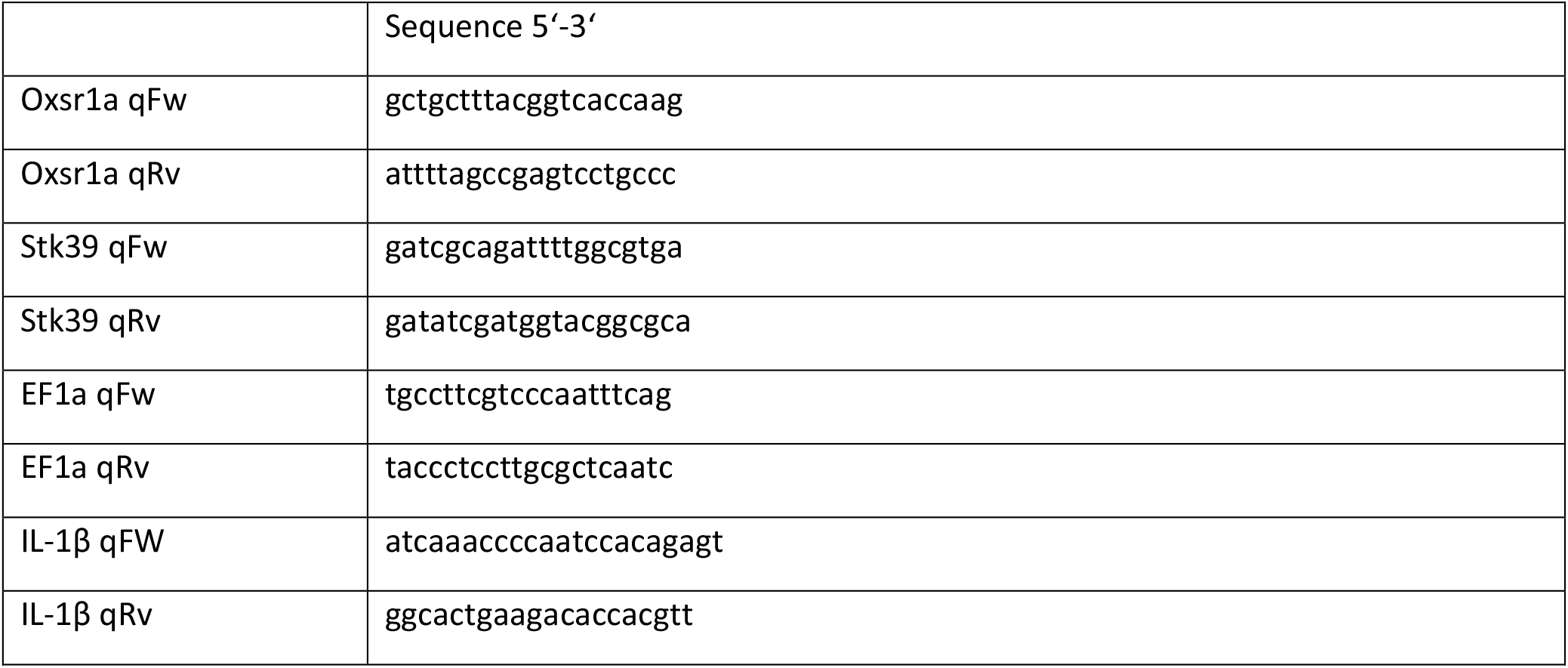
Primers used for gene expression studies.

**Table 2.**
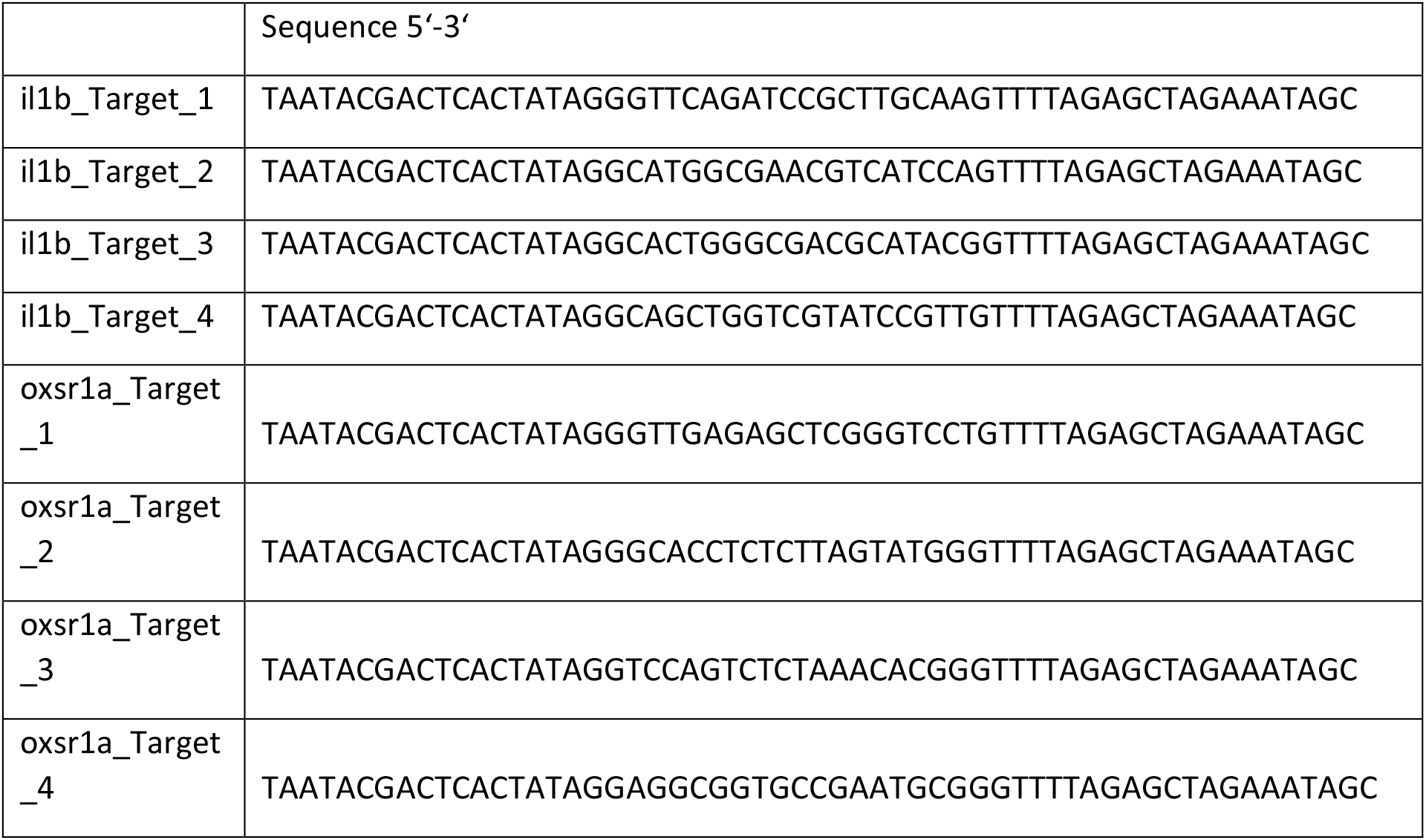
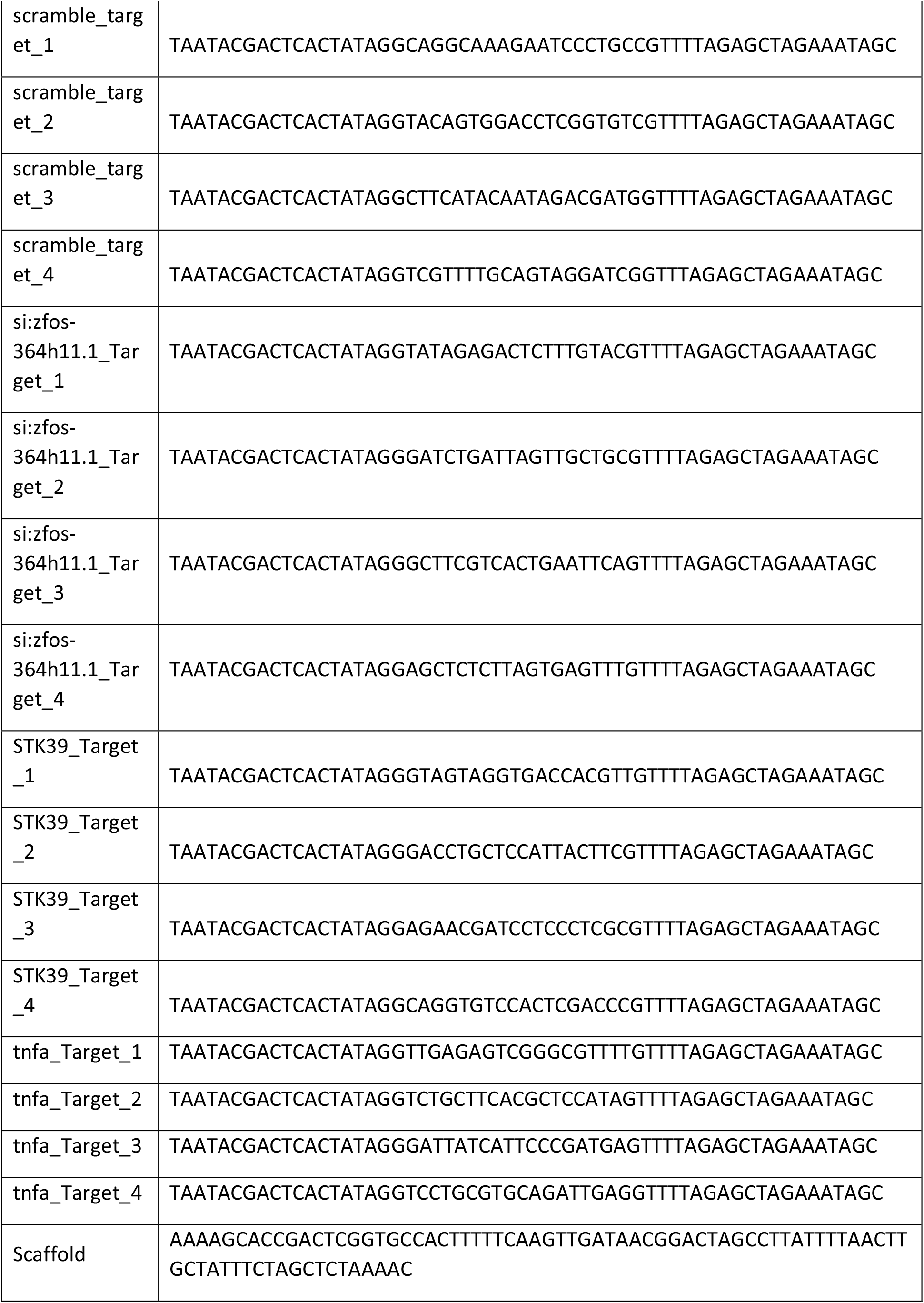
Primers used to generate gRNAs for CRISPR-Cas9 knockdown.

**Table 3.**
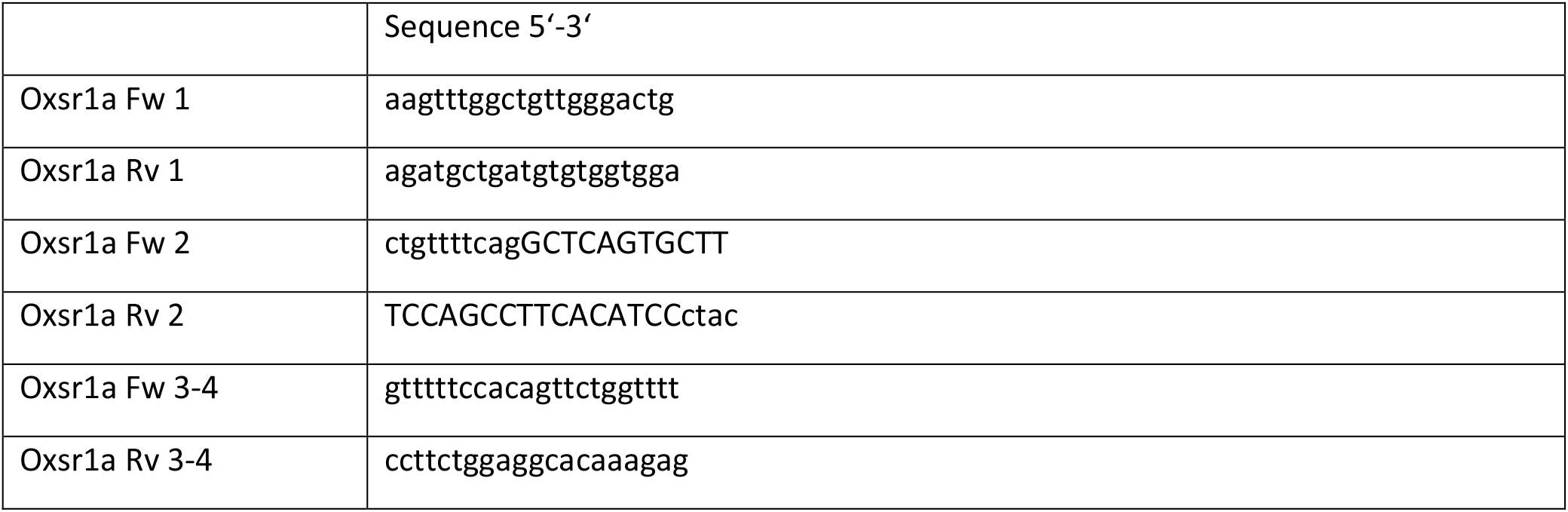
Primers used for genotyping the *oxsr1a*^*syd5*^ allele.

**Table 4.** Survival data of zebrafish embryos treated at 1 day post fertilization with varying concentrations of Compound B (CB). Drug was administered once at the beginning of the experiment.

| CB (μM) | % Survival |  |  |
| --- | --- | --- | --- |
|  | 24hrs | 48hrs | 120hrs |
| 0.9 | 100 | 100 | 100 |
| 1.8 | 100 | 100 | 100 |
| 3.6 | 100 | 100 | 47 |
| 7.5 | 100 | 47 | 0 |
| 15 | 5.8 | 0 | 0 |
| 30 | 0 | 0 | 0 |

**Figure 3:**
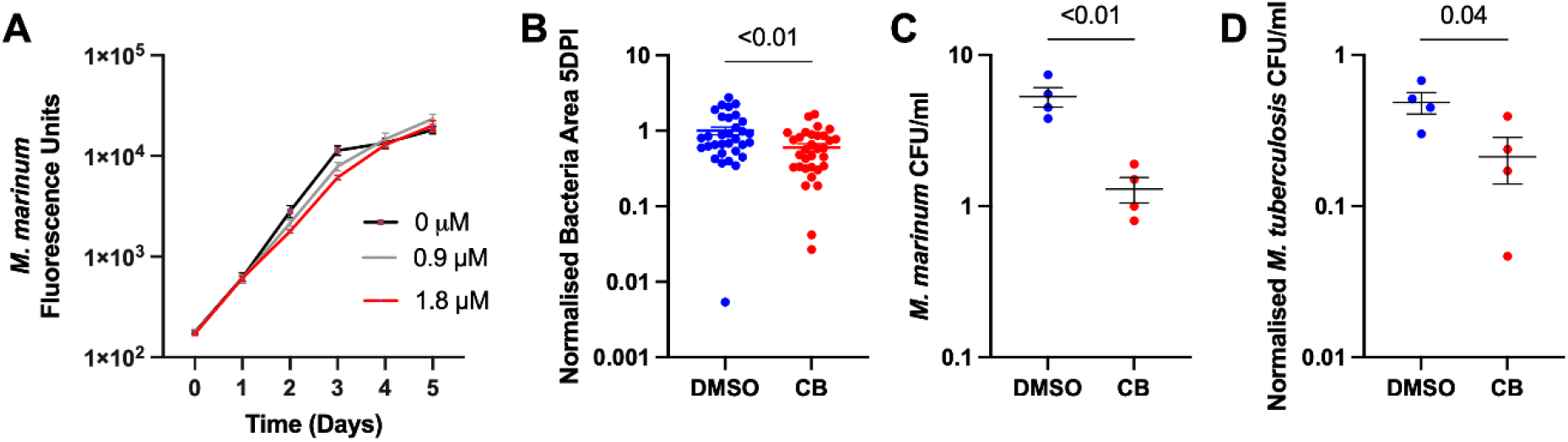
Compound B inhibition of SPAK/OXSR1 reduces mycobacterial burden in zebrafish embryos and human THP-1 cells. A) Quantification of axenic *M*. marinum-tdTomato growth by fluorescence in 7H9 broth culture supplemented with Compound B (CB). Red line (1.8 μM) indicates the concentration used to treat infected embryos. B) Quantification of *M. marinum* bacterial burden in CB-treated zebrafish embryos. C) Quantification of intracellular *M. marinum* burden in 1 dpi CB-treated PMA-differentiated THP-1 cells. D) Quantification of intracellular *M. tuberculosis* burden in 3 dpi CB-treated PMA-differentiated THP-1 cells.

Immersion of *M. marinum-*infected zebrafish embryos in 1.8 μM CB immediately after infection replicated the effect of *oxsr1a* knockdown by decreasing bacterial burden (Figure 3B). Treatment of infected THP-1 cells with CB also phenocopied the effect of *OXSR1* knockdown, with reduced *M. marinum* burden at 1 dpi (Figure 3C) and reduced *M. tuberculosis* H37Rv burden at 3 dpi compared to DMSO treatment (Figure 3D).

### The *M. marinum* ESX1 secretion system is required for infection-induced upregulation of host *oxsr1a*

We next examined the role of mycobacterial virulence in driving infection-induced expression of *oxsr1a* and sensitivity to Oxsr1a depletion or inhibition by infecting zebrafish embryos with ΔESX1 *M. marinum*, which cannot escape the macrophage phagocytic vacuole and fails to activate the inflammasome^31^. In contrast to infections with WT *M. marinum*, we did not observe an upregulation of either kinase when zebrafish embryos were infected with ΔESX1 *M. marinum* (Figure 4A). The burdens of zebrafish embryos infected with ΔESX1 *M. marinum* were unsensitive *oxsr1a* knockdown (Figure 4B) or treatment with CB (Figure 4C) suggesting a potential role for inflammasome activation in the protective effects of Oxsr1a depletion or inhibition.

**Figure 4:**
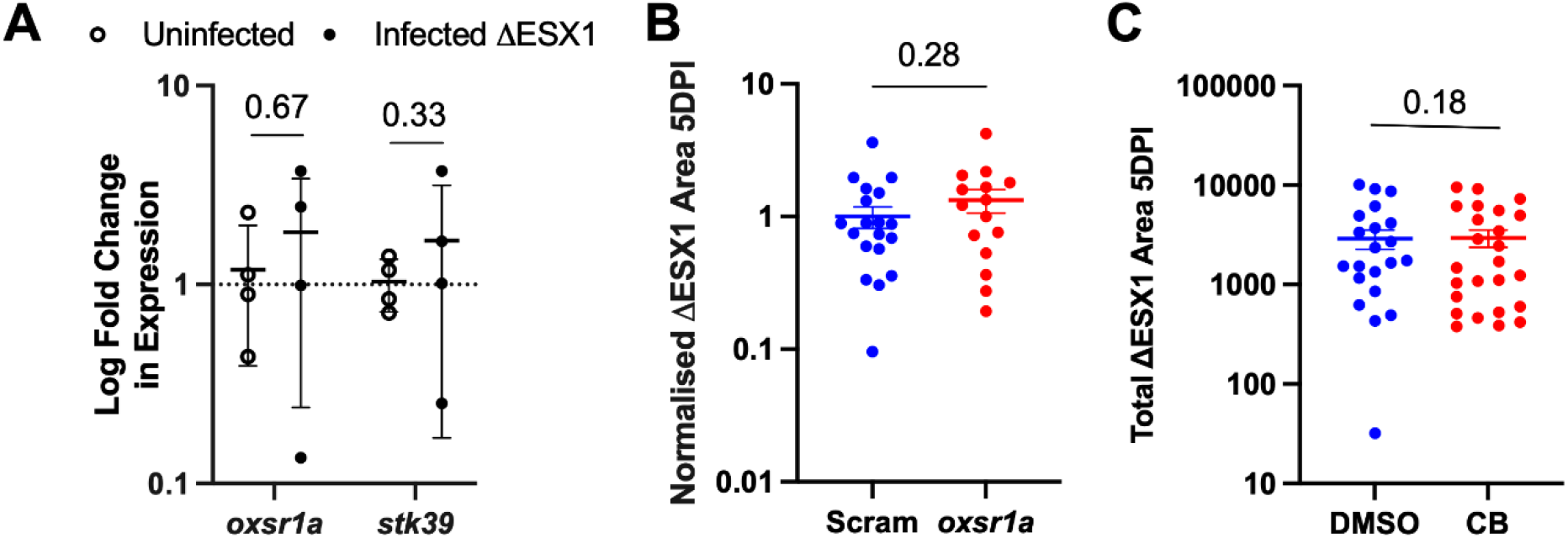
*M. marinum* ESX1 drives infection-induced *oxsr1a* expression. A) Relative expression of *oxsr1a* and *stk39* in zebrafish embryos at 3 days post infection with ΔESX1-*M. marinum* compared to age-matched uninfected controls. Biological replicates (n=4) represent pooled RNA from 7-10 embryos. B) Quantification of ΔESX1-*M. marinum* bacterial burden in scramble control and mosaic F0 *oxsr1a* CRISPR embryos. C) Quantification of ΔESX1-*M. marinum* bacterial burden in DMSO vehicle control and Compound B-treated embryos.

### Infection-induced OXSR1 suppresses inflammasome activity to aid mycobacterial infection

To determine if the reduced bacterial burden in *oxsr1a* knockdown was mediated by increased inflammasome activation, we used CRISPR to knockdown *si:zfos-364h11*.*1*, a zebrafish protein with orthology to mouse and rat NLRP3, hereafter referred to as *nlrp3*, and the *il1b* gene which encodes IL-1β (Extended Data 1). Knockdown of *nlrp3* alone did not affect the *M. marinum* burden but ameliorated the protective effect of *oxsr1a* knockdown against *M. marinum* infection (Figures 5A and 5B). The same effect was observed in zebrafish embryos subjected to *il1b* knockdown in combination with *oxsr1a* knockdown during *M. marinum* infection (Figure 5C).

**Figure 5:**
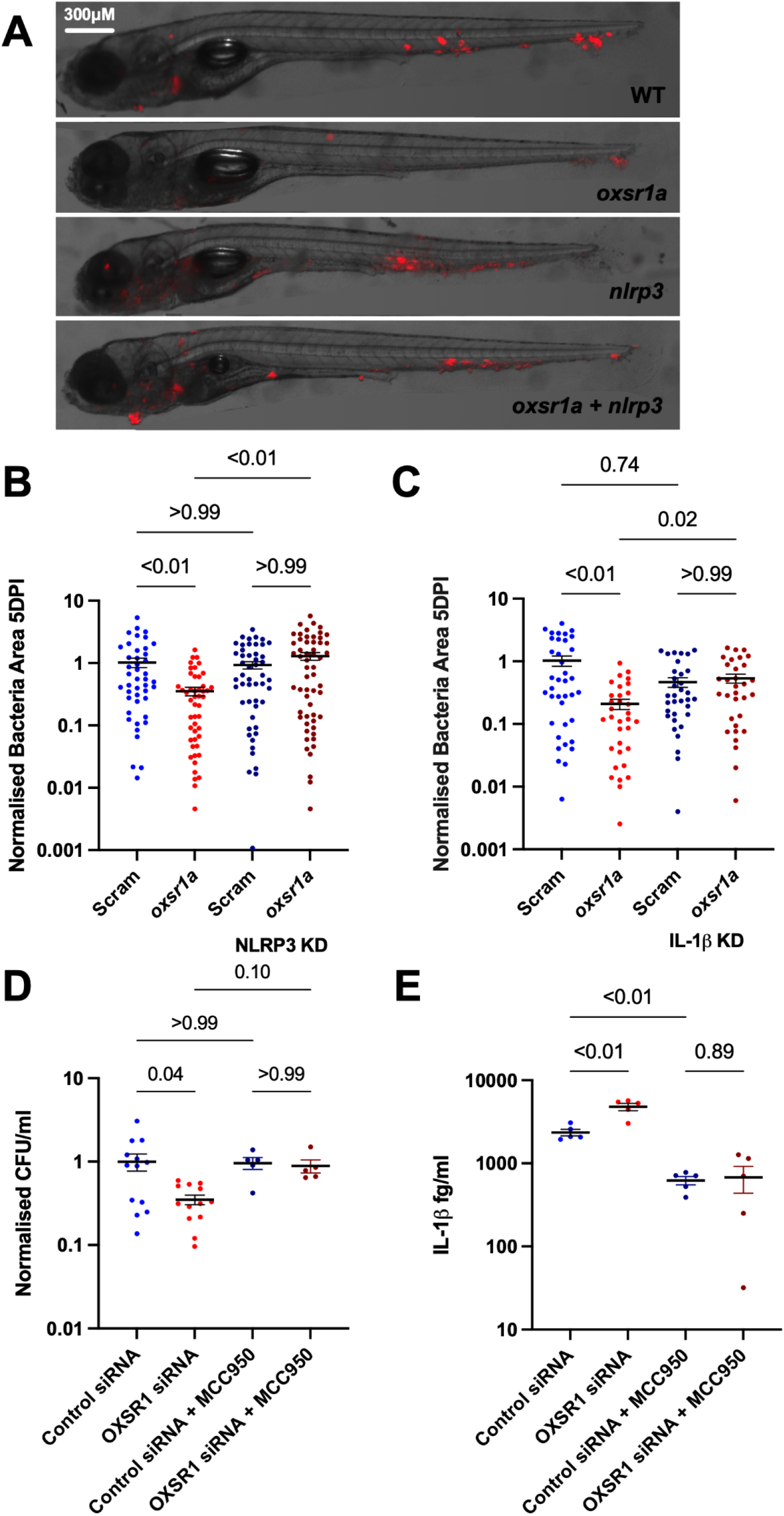
Infection-induced OXSR1 suppresses inflammasome activity to aid mycobacterial infection. A) Representative images of *M. marinum*-tdTomato (red) bacterial burden in 5 dpi WT, mosaic F0 *oxsr1a, nlrp3*, and dual *oxsr1a nlrp3* crispant embryos. Scale bar represents 300 µm. B) Quantification of *M. marinum* bacterial burden in scramble control, mosaic F0 *oxsr1a, nlrp3*, and dual *oxsr1a nlrp3* crispant embryos. Combined results of 3 biological replicates. C) Quantification of *M. marinum* bacterial burden in scramble control, mosaic F0 *oxsr1a, Il1b*, and dual *oxsr1a Il1b* crispant embryos. Combined results of 2 biological replicates. D) Quantification of intracellular *M. tuberculosis* bacterial burden in 3 dpi O*XSR1* knockdown PMA-differentiated THP-1 cells. E) IL-1β content in the supernatant of 3 dpi *M. tuberculosis*-infected O*XSR1* knockdown PMA-differentiated THP-1 cells.

In THP-1 cells infected with *M. tuberculosis* H37Rv, the *OXSR1* knockdown-mediated reduction in *M. tuberculosis* CFU observed at 3 dpi was ameliorated by treatment with the NLRP3 inhibitor MCC950 (Figure 5D). Supernatant IL-1β was significantly higher in media from *OXSR1* knockdown cells compared to the WT THP-1 cells after *M. tuberculosis* H37Rv infection and this increase in IL-1β was ablated by MCC950 treatment (Figure 5E). Together these data indicate that infection-induced increased expression of *oxsr1a* increases the mycobacterial burden through suppression of inflammasome activation.

### Infection-induced OXSR1 suppresses host protective TNF-α and cell death early in infection

Inflammasome-mediated IL-1β increases the macrophage killing of mycobacteria through upregulation of TNF-α^2^. We therefore repeated our infection experiments in *TgBAC(tnfa:GFP)*^*pd1028*^ embryos to determine if increased TNF-α production was mediating the resistance to mycobacterial infection in *oxsr1a* knockdown zebrafish. The ratio of *tnfa* promoter activity driven GFP per mycobacteria was increased specifically at sites of infection in *oxsr1a* knockdown embryos (Figure 6A) and also in Compound B-treated embryos (Figure 6B). This effect was dependent on *il1b* expression as knockdown of *il1b* suppressed *TgBAC(tnfa:GFP)*^*pd1028*^-driven GFP expression around sites of infection (Figure 6C).

**Figure 6:**
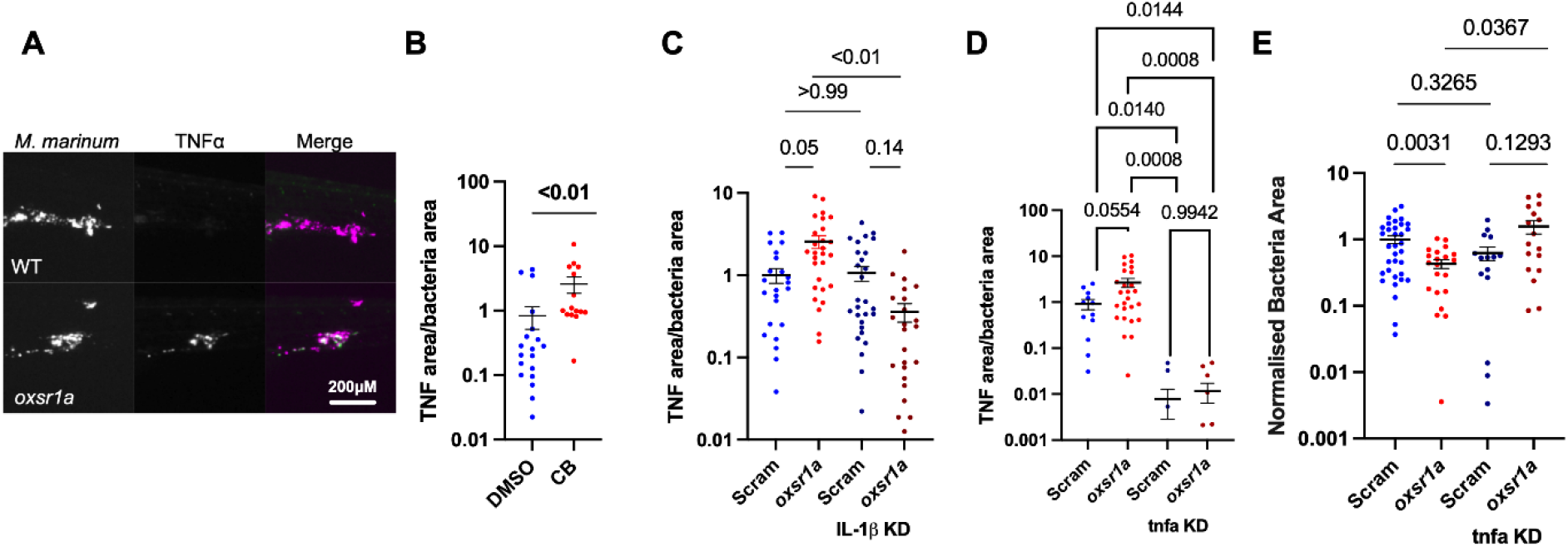
Infection-induced OXSR1 suppresses inflammasome activity to aid mycobacterial infection. A) Representative images of *Tg(tnfa:GFP)*^*pd1028*^ fluorescence around *M. marinum* granulomas. Scale bar represents 200 µm. B) Quantification of *Tg(tnfa:GFP)*^*pd1028*^ fluorescent pixels per bacterial pixel in *M. marinum*-infected CB-treated zebrafish embryos. C) Quantification of *Tg(tnfa:GFP)*^*pd1028*^ fluorescent pixels per bacterial pixel in *M. marinum*-infected WT, mosaic F0 *oxsr1a, Il1b*, and dual *oxsr1a Il1b* crispant embryos. Combined results of 2 independent experiments. D) Quantification of *Tg(tnfa:GFP)*^*pd1028*^ fluorescent pixels per bacterial pixel in *M. marinum*-infected WT, mosaic F0 *oxsr1a, tnfa*, and dual *oxsr1a tnfa* crispant embryos. Combined results of 2 independent experiments. E) Quantification of *M. marinum* burden in WT, mosaic F0 *oxsr1a, tnfa*, and dual *oxsr1a tnfa* crispant embryos. Combined results of 2 independent experiments.

To determine if *tnfa* expression acts downstream of *oxsr1a* depletion, we knocked down *tnfa* expression with CRISPR-Cas9 in the *TgBAC(tnfa:GFP)*^*pd1028*^ background to monitor knockdown efficacy (Figure 6D) ^32^. Knockdown of *tnfa* reduced the amount of infection-induced *tnfa* promoter-driven GFP produced around sites of infection and ameliorated the protective effect of *oxsr1a* knockdown against *M. marinum* infection (Figure 6E). Together these data suggest increased TNF-α downstream of inflammasome-processed IL-1β is the mechanism driving the lower bacterial burden in *oxsr1a* knockdown embryos.

## Discussion

Here we used the zebrafish-*M. marinum* and *in vitro* human-*M. tuberculosis* experimental systems to show that the WNK-OSXR1 signalling pathway has a critical role in infection-induced activation of the inflammasome. We present evidence that pathogenic mycobacteria increase macrophage K^+^ concentration by inducing expression of OXSR1. Infection-induced OXSR1 suppresses protective NLRP3 inflammasome responses and downstream IL-1β/ TNF-α production. Several studies have suggested that mycobacteria modulate inflammasome activation either by active inhibition or by upregulation of NOS, IFN-β and other negative inflammasome regulators^10^. Our data expand this literature by showing that mycobacterial infection-induced OXSR1 expression reduces protective inflammasome activation.

In our infection model we found that WT, but not a-virulent ΔESX1, *M. marinum* induced expression of both OXSR1 and SPAK. The ESX1 secretion system is essential for the virulence of *M. marinum* and is required for escape of the mycobacteria into the cytoplasm ^33,34^. This suggests that SPAK/OXSR1 upregulation is driven by the bacteria and fits with the well-established paradigm that pathogenic mycobacteria co-opt host pathways to establish persistent infection^35-39^.

We found *stk39* knockdown had no effect on host control of mycobacterial infection in zebrafish embryos. Previous studies have shown that mouse SPAK can play a role in activating macrophage inflammation in both lung injury and inflammatory bowel disease models^40-42^. These data raise the possibility that SPAK and OXSR1 may have species or organ specific roles in innate immunity and may respond differently to sterile and infectious triggers of inflammation.

The effect of Compound B on mycobacterial growth showed that small-molecule inhibition of OXSR1 can reproduce the impact of OXSR1 knockdown on mycobacterial survival. This observation provides proof of concept that OXSR1 may be a suitable target for host-directed therapies against mycobacterial and other intracellular infections. The reduction in bacterial burden was not as large in CB-treated fish as the reduction observed in OXSR1 knockdown embryos. This result may be because the maximum tolerated dose of CB was 1.8 µM, which is low compared to some reported EC_50_ values^30^. Therefore, CB may not have reduced OXSR1 activity to the same extent as in the OXSR1 knockdown embryos.

The results from the THP-1-derived macrophages confirm that the role of OXSR1 in the host response to infection is conserved across species. We showed that infection with WT *M. marinum* decreased the K^+^ content of OXSR1 knockdown cells but did not significantly affect the K^+^ content of WT cells. Paired with our data showing that WT *M. marinum* increases expression of *oxsr1a*, this suggests that virulent mycobacteria manipulate the SPAK/OXSR1 pathway to maintain high intracellular K^+^.

Several studies have shown that K^+^ concentration can affect mycobacterial growth and dormancy, and that successful colonisation of macrophages relies on the ability of the bacteria to maintain K^+^ homeostasis^43-45^. While here we have examined the effects of K^+^ efflux on inflammasome activation, it is possible that the high K^+^ maintained in WT cells also aids mycobacterial growth by helping the bacteria maintain the correct ion homeostasis. The mycobacterial infections in THP-1-derived macrophages revealed that both OXSR1 knockdown and treatment with CB resulted in reduced growth of both *M. marinum* and *M. tuberculosis* H37Rv in the human macrophage cell line. With *M. marinum* the maximum reduction was observed at 1 dpi, whereas with *M. tuberculosis* this was not seen until 3 days post infection. This is likely to be due to the differing replication times of both pathogens, which are 7 hours and 24 hours, respectively.

Here we have shown that OXSR1 knockdown can only reduce bacterial burden in zebrafish embryos if NLRP3, IL-1β, and TNFα are functional. In human cells we have shown that infected OXSR1 knockdown cells release significantly more IL-1β into the supernatant and that this is ablated by the NLRP3 inhibitor MCC950. Together this suggests that OXSR1 knockdown reduces bacterial burden via a first step of NLRP3 activation. Previous work in the zebrafish-*M. marinum* model has shown both host detrimental and host beneficial effects of inflammasome activation. While morpholino knockdown of *il1b* has been reported to increase bacterial burden, suggesting that *il1b* plays a host protective role; morpholino knockdown of *caspa* reduced bacterial burden, suggesting caspase-associated cell death of infected macrophages benefits the bacteria ^46^. In our experiments we did not find any effect of *il1b* or *nlrp3* knockdown on bacterial burden compared to control embryos. This may have been because we were using mosaic F0 CRISPR knockout, which is not a complete removal or because of *M. marinum* strain differences between studies.

We found either CB treatment and *oxsr1a* knockdown result in localised increased TNFα production at sites of infection. The fact that we only observed increased TNFα localised to sites of infection, and not throughout the whole embryo, suggests that *oxsr1a* knockdown primes cells for NLRP3 activation but does not cause excess systemic inflammation. Full activation of NLRP3 requires both a priming signal, to induce transcription of NLRP3 components, and pro-IL-1β and an activation signal, to induce oligomerization of NLRP3^18,19^. K^+^ efflux should provide only the second signal^18,23^; therefore in cells which have not been primed by infection with bacteria we would not expect to see significant NLRP3 activation. This suggests that OXSR1 inhibition may be an effective host-directed therapy strategy that induces beneficial inflammation at sites of infection without inducing detrimental systemic inflammation.

Our findings that OXSR1 can be targeted to decrease bacterial burden define a new avenue for the development of host-directed therapy. Although numerous studies have investigated the potential of inhibiting NLRP3 to minimize pathology ^47^, the possibility of activating inflammasomes to increase pathogen clearance has been largely unexplored. Here we have shown that enhancing inflammasome activation via K^+^ efflux can provide the dual benefits of maximising the anti-pathogen effects of inflammation without causing excess tissue damage. Given mycobacteria are not the only pathogens which inhibit inflammasome activation, OXSR1 inhibition may be an effective host-directed therapy with broad applicability.

## Methods

### Zebrafish husbandry

Adult zebrafish were housed at the Centenary Institute (Sydney Local Health District AWC Approval 2017-036). Zebrafish embryos were obtained by natural spawning and embryos were raised at 28°C in E3 media.

### Zebrafish lines

Wild type zebrafish are the TAB background. Transgenic line was *Tg(tnfa:GFP)*^*pd1028* 48^.

### Infection of zebrafish embryos

Embryos were infected by microinjection with ∼400 fluorescent *M. marinum* M strain and ΔESX1 *M. marinum* as previously described^49^. Embryos were recovered into E3 supplemented with 0.036 g/L PTU, housed at 28 °C and imaged on day 5 of infection unless otherwise stated.

### Quantitative Reverse Transcription PCR (qRT-PCR)

RNA was extracted from 5-10 embryos using TRIzol (Invitrogen) according to the manufacturer’s instructions. Equal amounts of RNA (either 1 or 2 μg depending on RNA yield) were used for the cDNA synthesis reaction. qRT-PCR reactions were carried out on a Biorad CFX machine using ThermoFisher PowerUP SYBR green and primers described in Table 1. The relative quantity of transcripts was calculated by the Delta-delta CT method.

### Imaging

Live zebrafish embryos were anaesthetized in M-222 (Tricaine) and mounted in 3% methylcellulose for static imaging on a Leica M205FA fluorescence stereomicroscope. Fluorescent pixel count analyses were carried out with Image J Software Version 1.51j and intensity measurements were performed as previously described^49^.

### CRISPR-Cas9 knockdown and mutant generation

Primers used for gRNA transcription are detailed in Table 2 and were designed by Wu et al. ^50^. Templates for gRNA transcription were produced by annealing and amplifying gene specific oligos to the scaffold oligo using the NEB Q5 polymerase. Pooled transcription of gRNAs was carried out using the NEB HiScribe T7 High Yield RNA Synthesis Kit.

Embryos were injected at the single cell stage with an injection mix containing 1 μl phenol red, 2 μl 500 ng/μl pooled guides, and 2 μl of 10 μM Cas9. All ‘Scram’ embryos are injected with scrambled guide RNA.

To create oxsr1a knockout line, F0 crispants were outcrossed to WT AB, and HRM analysis was conducted on F1 progeny. F1s with a visible HRM shift using primers amplifying the 4 predicted cut sites (primer 3 spanned 2 cut sites) were sent for sanger sequencing. An F1 was discovered carrying an 8 bp deletion causing a premature stop at amino acid 13 (Extended Data 2). F2 progeny were genotyped with a custom KASP assay ordered from LGC Biosearch Technologies.

### Drug treatments

Embryos and cells were treated with vehicle control (DMSO or water as appropriate), 10 µM MCC950, 77 µM Ac-YVAD-cmk, 1.8 µM Compound B, or 48 µM Furosemide (Sigma) immediately after infection. For zebrafish, the drugs and E3 were replaced on days 0, 2, and 4 dpi. For cell culture, drugs were replaced at 4 hours post infection.

### Axenic culture

A mid-log culture of fluorescent *M. marinum* was diluted 1:100 and aliquoted into 96 well plates for drug treatment. Cultures were maintained at 28°C in a static incubator and bacterial fluorescence was measured in a BMG Fluorostar plate reader.

### THP-1 cell culture

Human THP-1 cells (ATCC^®^ TIB-202^™^) were cultured in RPMI media (22400089, ThermoFisher) supplemented with 1% (v/v) non-essential amino acids (11140050, ThermoFisher), 1 mM sodium pyruvate (11360070, ThermoFisher), 10% (v/v) FCS (Hyclone, GE Healthcare) and 0.1 mg/ml penicillin/streptomycin (15140122, ThermoFisher) at 37°C, 5% CO_2_.

### Viral production

24 hours prior to transfection, 4×10^6^ HEK2937 cells were seeded in a 100 mm culture dish. On the day of transfection, cells were co-transfected with 15ug of the pLKO.1_GFP (#30323, Addgene) vector containing OXSR1_Sh1, OXSR1_Sh2 or AthmiR, 6.5 µg of the packaging plasmid pMDL-g/prre (#12251, Addgene), 2.5 µg of the packaging plasmid pRSV-Rev (#12253, Addgene) and 3.5 µg of the envelop expressing plasmid pMD2-VSV-G (#12259, Addgene) by the calcium phosphate transfection method. Culture media was changed the following day and cells were cultured for another 24 hours. Medium containing lentiviral particles was then collected, debris was cleared by centrifugation at 430 g for 5 minutes, filtered through a 0.45 µm filter, aliquoted, and stored at -80°C.

### Transduction

Briefly, 5×10^5^ THP-1 cells were resuspended in 500 µl of fresh culture media containing 10 µg/ml polybrene. After adding 50 µl of virus, cells were spinoculated for 90 min at 462 g, 22°C. After spinning, pelleted cells were resuspended in in the same media and incubated for 4 hours at 37C, 5%CO2. Following incubation, cells were pelleted, resuspended in fresh culture media and transferred to a 6 well plate. Cells were cultured for 48 hours before FACS selection.

### Western Blotting

Protein lysates were loaded onto 4-12% BIS-Tris Protein gels (NP0336BOX, ThermoFisher) for electrophoresis followed by transfer onto a PVDF membrane (MILIPVH00010, Merck Millipore). Membrane was blocked with 5% (v/v) skim milk for 1 hour at room temperature, incubated overnight with a 1:1000 dilution of rabbit anti-OXSR1 (ab97694, Abcam) followed by incubation with a 1:5000 dilution of a donkey anti-rabbit IgG HRP antibody (AP182P, Merck Millipore) and 1:5000 dilution of mouse anti-GAPDH (ab8245, Abcam) antibody, followed by incubation with a 1:5000 dilution of a donkey anti-mouse IgG HRP antibody (AP192P, Merck Millipore). Protein detection was performed using SuperSignal West Pico PLUS (34579, ThermoFisher) and imaged on a Bio-Rad ChemiDoc Imaging System.

### ION K+ Green (undifferentiated cells)

For flow cytometry, 2×10^5^ undifferentiated THP-1 cells/well (ATCC^®^ TIB-202^™^) were seeded into a 96 well plate and incubated at 37°C for 1.5-2 hrs with either Furosemide, 37.5 mM or 75 mM KCl. ION K+ Green was added to a final concentration of 52.8 mM and cells were incubated for a further 15 minutes. Cells were spun down for 5 minutes at 462 g and resuspended in PBS + 2% FCS supplemented with either Furosemide, 37.5 mM or 75 mM KCl. ION K+ Green fluorescence was captured on a BD Fortessa through the PE channel. 5000 events were captured per sample.

### ION K+ Green (differentiated cells)

2×10^5^ THP-1 cells/well were seeded into a 96 well plate and differentiated for 24 hrs with 100 mM PMA. Cells were then infected with frozen single cell preparation *M. marinum-*katushka at an MOI of 1. After 4 hrs extracellular bacteria were removed by washing with PBS + 2% FCS, and cells were incubated at 32°C for 3 days. Cells were lifted from the plate by 15 minute incubation at 37°C with Accutase™ (StemCell Technologies), then washed with PBS + 2% FCS. ION K+ Green was added to a final concentration of 52.8 mM and cells were incubated for a further 15 minutes. Cells were spun down for 5 minutes at 462 g and resuspended in PBS + 2% FCS for flow cytometry. ION K+ Green fluorescence was captured on a BD Fortessa through the FITC channel (so as not to overlap with katushka). 5000 events were captured per sample.

For static imaging, THP-1 cells were seeded onto an imaging slide coated with 1% low melting point agarose and differentiated for 24 hrs with 100 mM PMA. ION K+ Green was added to a final concentration of 52.8 mM and cells were incubated for 15 minutes at 37°C. Culture media was replaced with PBS + 2% FCS. Cells were imaged on a Leica Sp8 and mean ION K+ Green fluorescence was analysed using the ‘measure’ function in Image J.

### Mycobacterial infection of THP-1 cells

2⨯10^5^ THP-1 cells/well were seeded into a 96 well plate and differentiated for 24 hrs with 100 mM PMA. Cells were then infected with either mid log culture of *M. tuberculosis* H37Rv or frozen single cell preparation of *M. marinum* at an MOI of 1. After 4 hrs, extracellular bacteria were removed by washing with PBS + 2% FCS, and cells were incubated at either 32°C for *M. marinum* infections or 37°C for *M. tuberculosis* infections.

### Mycobacterial CFU recovery from THP-1 cells

Cells were washed in PBS + 2% FCS and lysed with TDW + 1% Triton X100 for 10 minutes. lysate was serially diluted and plated on 7H10 agar supplemented with 50 μg/ml hygromycin for the recovery of *M. marinum* or a mix of 200,000 units/L polymyxin B, 50mg/L carbenicillin, 10mg/L amphotericin B, and 20mg/L trimethoprim lactate for the recovery of *M. tuberculosis*. Plates were incubated at 32°C for 7 days (*M. marinum*) or 37°C for 14 days (*M. tuberculosis)*.

### Measurement of human IL-1βin supernatants

IL-1β was measured by cytometric bead array, using a human IL-1β enhanced-sensitivity flex set (BD Biosciences). Undiluted cell supernatant was stained according to the manufacturer’s instructions and run on a BD FACS Canto II. Data was analysed using FCAP array software.

### Statistics

All statistical tests were calculated in Graphpad Prism. T-tests were unpaired t-tests with Welch’s correction. All ANOVA were ordinary one-way ANOVA, comparing the means of specified pairings, using Turkey’s multiple comparisons test with a single pooled variance. In cases where data was pooled from multiple experiments, data from each was normalized to its own within-experiment control (usually DMSO) before pooling. Error bars indicate SEM. Outliers were removed using ROUT, with Q=1%.

## Acknowledgements

We thank Dr Kristina Jahn of Sydney Cytometry for assistance with imaging equipment; Ms Kaiming Luo and Dr Pradeep Cholan for technical assistance, and all members of the Tuberculosis Research Program at the Centenary Institute for helpful comments.

## Author contributions

E.H. and S.H.O designed the experiments. E.H., V.L.T, S.H.O. performed the experiments. A.R.M.F. performed microscopy. N.P. and J.J-L.W. generated knockdown cell lines. E.H. and S.O. wrote the paper that was reviewed by all authors. W.J.B., and S.H.O. supervised the project.

---

**Extended Data 1.**
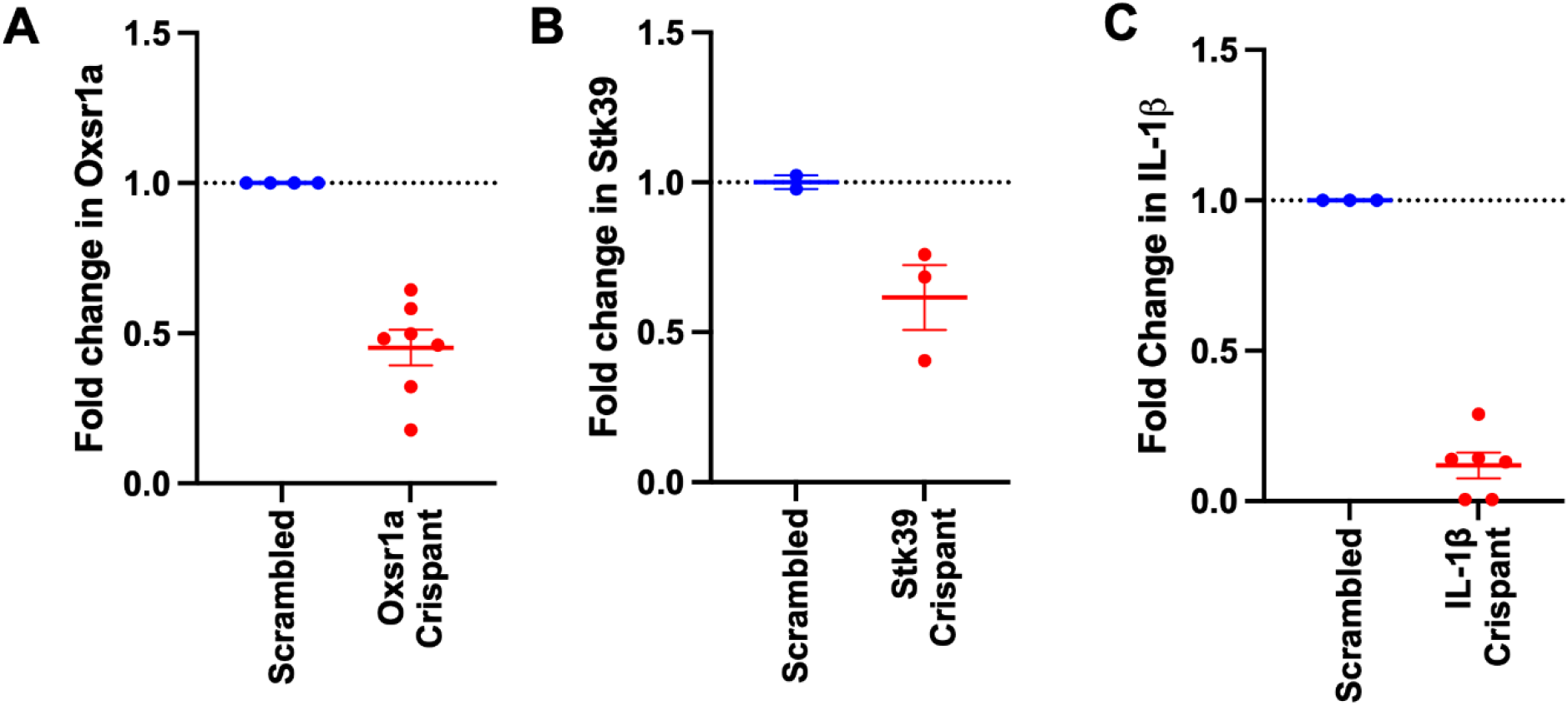
Knockdown efficacy for CRISPR-Cas9 depletion of A) *oxsr1a*, B) *stk39*, C) *il1b*, in zebrafish embryos.

**Extended Data 2.**
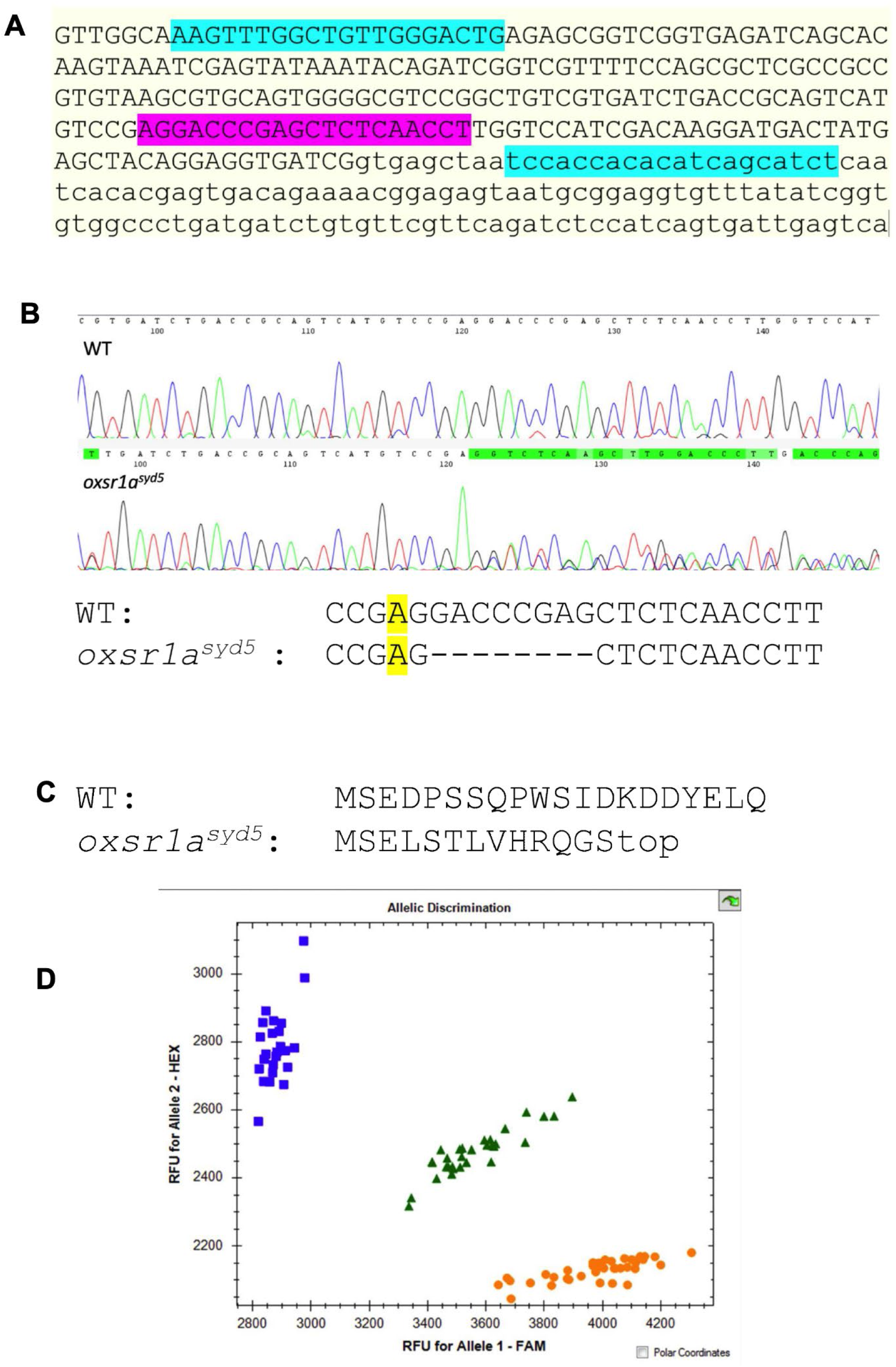
Generation of the *oxsr1a*^*syd5*^ allele. A) Map of primer binding sites (blue) and gRNA binding site (pink) around the *oxsr1a*^*syd5*^ allele. Sequence shows exon 1 (upper case) and part of intron 1 (lower case) of *oxsr1a* B) Chromatograms showing partial sequence of the *oxsr1a*^*syd5*^ allele. C) Predicted OXSR1 protein sequence of the *oxsr1a*^*syd5*^ allele. D) Representative genotyping results of hetxhet F2 cross. FAM = WT allele; HEX = *oxsr1a*^*syd5*^ allele.

**Extended Data 3.**
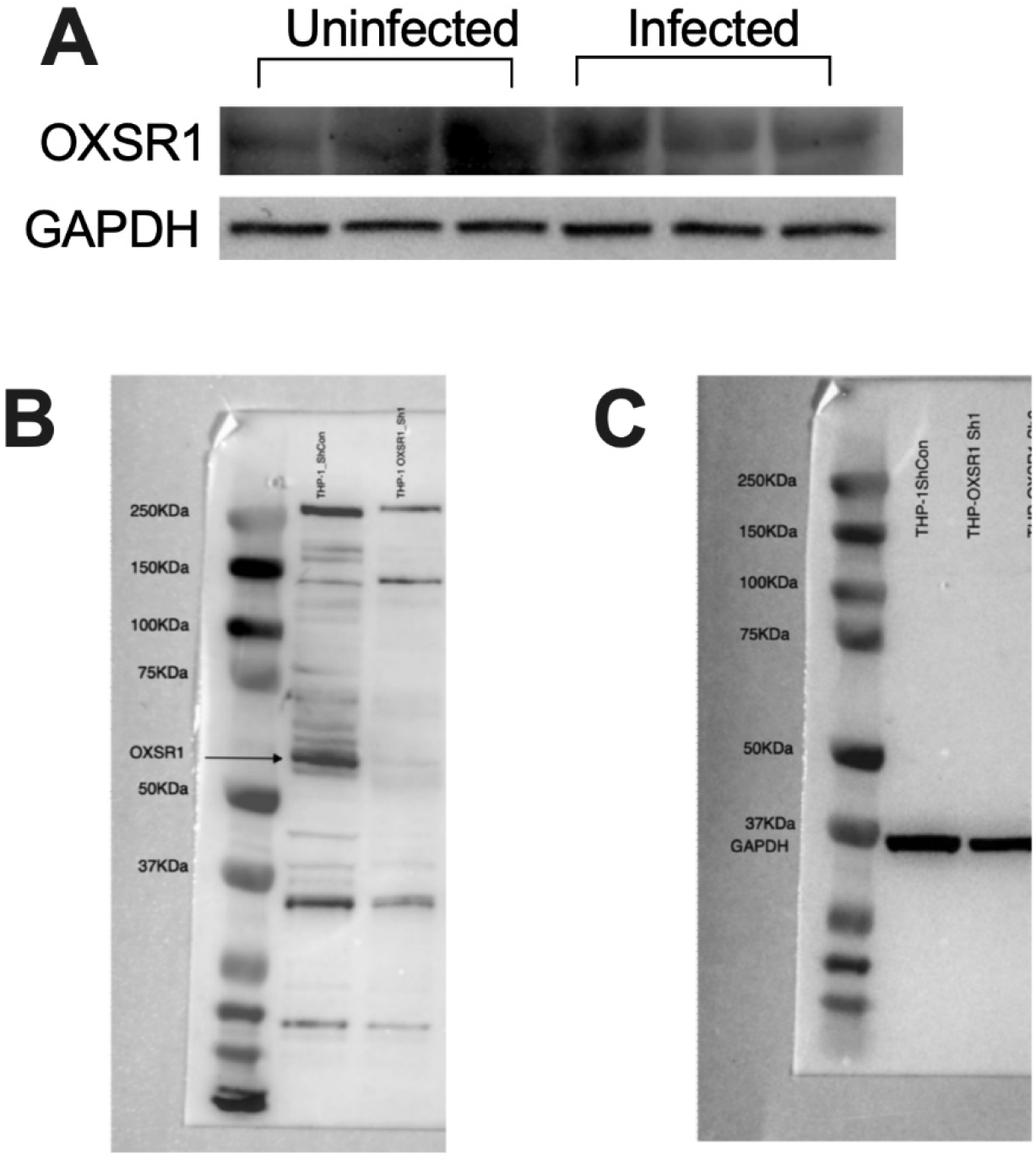
A) Western blot of OXSR1 and GAPDH protein levels in 3 dpi PMA-differentiated *M. tuberculosis*-infected THP-1 cells. Full western blots showing OXSR1 (B) and GAPDH (C) protein levels in THP-1 control and knockdown cell lines.

